# Guanosine tetraphosphate (ppGpp) accumulation inhibits chloroplast gene expression and promotes super grana formation in the moss *Physcomitrium* (*Physcomitrella*) *patens*

**DOI:** 10.1101/2021.01.06.425534

**Authors:** Seddik Harchouni, Samantha England, Julien Vieu, Aicha Aouane, Sylvie Citerne, Bertrand Legeret, Yonghua Li-Beisson, Benoît Menand, Ben Field

## Abstract

The nucleotides guanosine tetraphosphate and pentaphosphate (or ppGpp) are implicated in the regulation of chloroplast function in plants. ppGpp signalling is best understood in the model vascular plant *Arabidopsis thaliana* where it acts to regulate plastid gene expression to influence photosynthesis, plant development and immunity. However, little is known about the conservation or diversity of ppGpp signaling in other land plants. Here, we studied the function of ppGpp in the moss *Physcomitrium* (previously *Physcomitrella*) *patens* using an inducible system for triggering ppGpp accumulation. We used this approach to investigate the effects of ppGpp on chloroplast function, photosynthesis and growth. We demonstrate that ppGpp accumulation causes a dramatic drop in photosynthetic capacity by inhibiting chloroplast gene expression. This was accompanied by the unexpected reorganisation of the thylakoid system into super grana. Surprisingly, these changes did not affect gametophore growth, suggesting that bryophytes and vascular plants may have different tolerances to defects in photosynthesis. Our findings point to the existence of both highly conserved and more specific targets of ppGpp signalling in the land plants that may reflect different growth strategies.

## Introduction

Guanosine tetraphosphate and pentaphosphate (or ppGpp) are nucleotides that are implicated in stress acclimation in plants and bacteria. Bacterial ppGpp signalling is intensely investigated and mechanistically diverse (Hauryliuk *et al*., 2015; Steinchen & Bange, 2016; Ronneau & Hallez, 2019). Various stresses and environmental changes can trigger the synthesis of ppGpp from ATP and GDP/GTP by enzymes of the RelA SpoT Homologue (RSH) superfamily. The resulting increase in ppGpp concentration acts to reduce proliferation and activate acclimatory pathways by targeting enzymes involved in transcription, translation, metabolism and replication.

In plants, ppGpp signalling is less well characterised and its diversity is essentially unexplored. At least three conserved families of chloroplast-targeted RSH enzymes for both ppGpp synthesis and hydrolysis can be found in land plants. These are named RSH1, RSH2/3 and RSH4 (Atkinson *et al*., 2011; Ito *et al*., 2017; Avilan *et al*., 2019). In the angiosperm *Arabidopsis thaliana*, where ppGpp signalling is best characterised, ppGpp is known to act as a potent inhibitor of chloroplast gene expression *in vivo* (Maekawa, Mikika *et al*., 2015; Yamburenko *et al*., 2015; Sugliani *et al*., 2016). Increased ppGpp levels lead to major changes in the stoichiometry of photosynthetic complexes such as photosystem II (PSII), and cause a general inhibition of photosynthesis (Maekawa, M. *et al*., 2015; Sugliani *et al*., 2016; Honoki *et al*., 2018). *RSH* mutants with modified ppGpp content are affected in photosynthesis, development and immunity (Sugliani *et al*., 2016; Abdelkefi *et al*., 2018). ppGpp is also known to accumulate in response to various different abiotic stresses (Takahashi *et al*., 2004; Ihara *et al*., 2015), and when ectopically produced improves the tolerance of plants to nitrogen deprivation (Maekawa, M. *et al*., 2015; Honoki *et al*., 2018).

Relatively little is known about ppGpp signalling in other plants. One of the few studies outside of *A. thaliana* is on the function of RSH enzymes in the moss *Physcomitrium patens* (also known as *Physcomitrella patens* and hereafter named *P. patens*) (Sato *et al*., 2015; Rensing *et al*., 2020). As a representant of the non-vascular land plants (bryophytes), *P. patens* occupies an interesting phylogenetic position for the study of evolution of plant traits, particularly in comparison with vascular plants (Harrison & Morris, 2018). The *P. patens* genome contains nine *RSH* genes, encoding members of the RSH1, RSH2/3 and RSH4 families, that show different tissue and stress responsive expression patterns (Sato *et al*., 2015). *PpRSH2a* and *PpRSH2b* appear to play a major role in *P. patens* ppGpp biosynthesis because they are highly expressed and encode chloroplast targeted enzymes that show ppGpp synthetase activity *in vitro*. Overexpression of *PpRSH2a* or *PpRSH2b* resulted in a mild slow growth phenotype in *P. patens* grown on media supplemented with glucose (Sato *et al*., 2015). However, the developmental stage affected was not clear as the moss grown in this condition seemed to be a mix of filamentous protonema and leafy gametophores. Furthermore, comparisons with works on *A. thaliana* are limited because chloroplast phenotypes and photosynthetic capacity were not analysed and we do not know whether ppGpp levels increase in the *P. patens* RSH overexpression lines (Maekawa, M. *et al*., 2015; Sato *et al*., 2015; Sugliani *et al*., 2016). To address these problems and to take advantage of the informative evolutionary position of *P. patens* we developed a new strategy to investigate the role of ppGpp. We created a system for the inducible expression of a bacterial ppGpp synthetase that allowed us to increase ppGpp content in the chloroplast independently of endogenous RSH enzymes or their regulatory mechanisms, and in a precisely controllable manner. We focussed our study *on P. patens* shoots, called gametophores, because they are typically the most prominent vegetative developmental stage of moss in the wild. Gametophores also carry leaf-like structures called phyllids that evolved independently to the leaves of vascular plants and are particurly interesting for comparative biology (Fujita *et al*., 2008; Harrison & Morris, 2018; Glime, 2020; Rensing *et al*., 2020). In addition, phyllids are a single-cell-layer thick and are therefore highly convenient for microscopic observation (Rensing *et al*., 2020). Using this approach, we discovered that ppGpp accumulation caused a dramatic drop in photosynthetic capacity by inhibiting chloroplast gene expression that was accompanied by an unexpected reorganisation of the thylakoid system into super grana. Surprisingly, these changes did not appear to significantly affect growth or development. Our results shed new light on the mechanisms and conservation of ppGpp signalling in land plants.

## Results

### An inducible expression system for ppGpp accumulation in *P. patens*

To study the role of ppGpp in *P. patens* we created inducible expression lines that upon treatment with estradiol activate the expression of a gene encoding the ppGpp synthase domain of *E. coli* RelA fused to a chloroplast target peptide (SYN) (Fig. 1A). The inducible expression cassette bearing the SYN gene was inserted at the neutral PIG1 locus (*P. patens* InterGenic 1) (Okano *et al*., 2009) by gene targeting (Fig. S1A). In parallel, control lines (SYN^D>G^) were created using an inducible cassette encoding a catalytically inactivated version of the SYN gene. SYN and SYN^D>G^ lines with stable single insertions at the target locus were identified by Southern blotting and used for all subsequent experiments (Fig. S1B).

**Figure 1.**
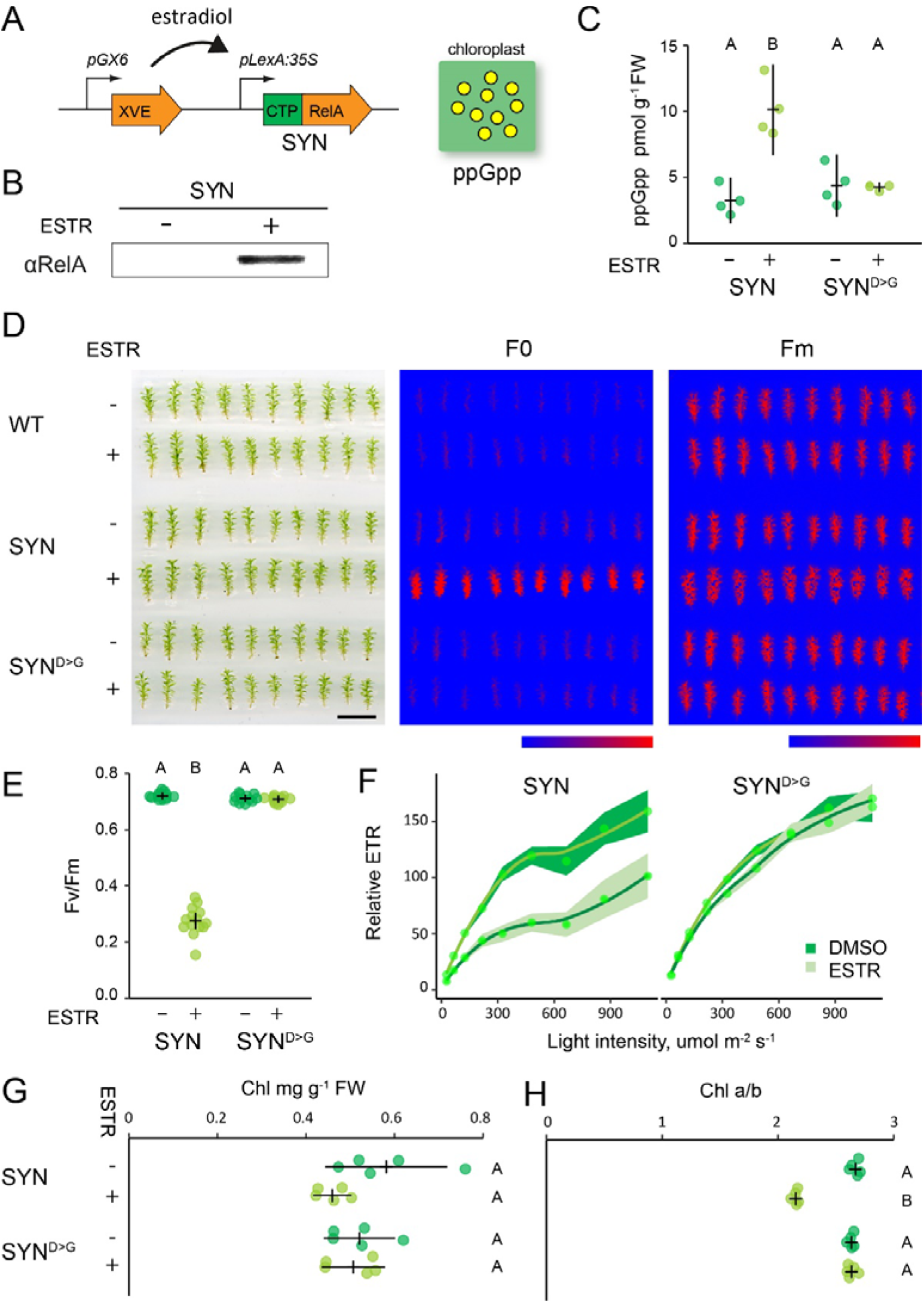
Inducible ppGpp production reduces photosynthetic activity. (A) Schematic representation of the estradiol inducible ppGpp overproduction system in SYN lines. (B) Immunoblot on total protein from SYN gametophores after 35 days of induction (DOI) with 0 μM (-) or 5 μM estradiol (+) using an anti-RelA antibody. (C) ppGpp levels in SYN and control SYN^D>G^ lines after 35 DOI (n=4 biological repeats). The photosynthetic status of SYN and SYN^D>G^ gametophores was investigated (D) by imaging basal (F0) and maximal (Fm) chlorophyll fluorescence (scale, 1 cm; false color scale, 20-2000 intensity units), (E) quantifying the maximal efficiency of PSII (Fv/Fm) (n=12-14 gametophores, one-way ANOVA, *P*<0.0001), and (F) calculating the relative electron transport rate (n=15 gametophores). (G) Chlorophyll concentration and (H) chlorophyll *a/b* ratios of SYN and SYN^D>G^ gametophores. Experiments performed on gametophores after 35 DOI. Error bars and intervals indicate 95% CI. Significance was tested with one-way ANOVA and post hoc Dunnett test, *P*<0.0001.

We next characterized the properties of the inducible expression system on *P. patens* grown on peat pellets, a condition that favours gametophore development. Induction of SYN gametophores by treatment with estradiol led to the appearance of a protein band that reacted with an antibody raised against RelA, and whose size is consistent with the expected size of the mature SYN protein (Fig. 1B). No band was detected in SYN gametophores treated with solvent only, indicating that the estradiol inducible system maintains tight control over the expression of SYN. Expression of the SYN protein was accompanied by an increase in ppGpp to levels three-fold higher than in the noninduced control or the induced SYN^D>G^ control line (Fig. 1C). These data demonstrate that SYN lines are a suitable tool for triggering the *in vivo* production of ppGpp.

### ppGpp accumulation reduces photosynthetic activity

Studies in *A. thaliana* show that ppGpp accumulation causes a reduction in photosynthetic capacity (Maekawa, M. *et al*., 2015; Sugliani *et al*., 2016; Honoki *et al*., 2018). We therefore analyzed the photosynthetic parameters of induced SYN gametophores to determine whether an increase in ppGpp levels *in vivo* can affect photosynthesis in *P. patens*. We found that the induction of SYN resulted in a large increase in basal chlorophyll fluorescence (F0) that was not observed in the SYN^D>G^ control line or the wild-type, indicating that this change is specific to the accumulation of ppGpp and not other aspects of the inducible expression system (Fig. 1D). Consistent with the increase in F0, induced SYN gametophores also showed a large drop in the maximal efficiency of PSII (Fv/Fm) (Fig 1E). The average Fv/Fm was 0.27 ± 0.013 (mean ± SE) in induced SYN gametophores and 0.70 ± 0.003 (mean ± SE) in induced SYN^D>G^ gametophores. (Fig. 1E). The decrease in Fv/Fm following SYN induction was detectable 2 days after induction and reached its lowest level after 35 days (Fig. S2). Similar changes in F0 and Fv/Fm were observed in all three independent SYN lines (Fig. S3). The reduction in Fv/Fm in SYN was also accompanied by a strong reduction in photosynthetic operating efficiency at all light intensities, indicating a reduced relative electron transfer rate (ETR) (Fig. 1F). Again, the reduced ETR was observed in all three SYN lines after induction but not in SYN^D>G^ (Fig. S4). While chlorophyll levels did not show clear evidence of a change in induced SYN gametophores (Fig. 1G), we observed a large and significant decrease in the chlorophyll *a*/*b* ratio, indicating major alterations in the organisation of the photosynthetic machinery (Fig. 1H). Surprisingly, despite the major defects in photosynthesis that we observed in induced SYN lines we did not observe a detectable effect on gametophore growth (Fig. 1D, S2). Altogether, these results indicate that ppGpp accumulation in SYN gametophores reduces photosynthetic capacity without affecting growth.

### ppGpp accumulation inhibits chloroplast gene expression

The alterations in photosynthetic capacity, chlorophyll fluorescence, and chlorophyll a/b ratio observed in induced SYN gametophores indicate that there is a fundamental remodelling of the photosynthetic machinery in response to ppGpp accumulation. To understand how the photosynthetic machinery is remodelled we analysed the expression of key photosynthetic proteins following SYN induction (Fig. 2A). First, we observed the progressive accumulation of the SYN protein and the SYN^D>G^ protein following induction of the respective lines. This was accompanied by the reduction in the amount of several chloroplast encoded photosynthesis proteins in the SYN line (Fig. 2A and Fig. S5, indicated in green). The most marked reduction was for RBCL, the large subunit of RUBISCO, and PsbA, a subunit of the PSII reaction centre (RC). These were followed by more modest reductions for CytF, a subunit of cytochrome B6f, and AtpB, a subunit of the ATPase. In contrast, we observed either no change or an increase in quantities of the nucleus encoded PSII and PSI light harvesting proteins (LHCB1, LHCB2, LHCA1), and the chloroplast encoded protein PsaA, a subunit of the PSI core. Interestingly, the unusual *P. patens* light harvesting protein LHCB9 also appeared to increase in the SYN line (Fig. 2A). This increase was even more pronounced in another experiment (Fig. S5). Together, these results indicate that ppGpp accumulation inhibits the accumulation of some but not all chloroplast encoded proteins.

**Figure 2.**
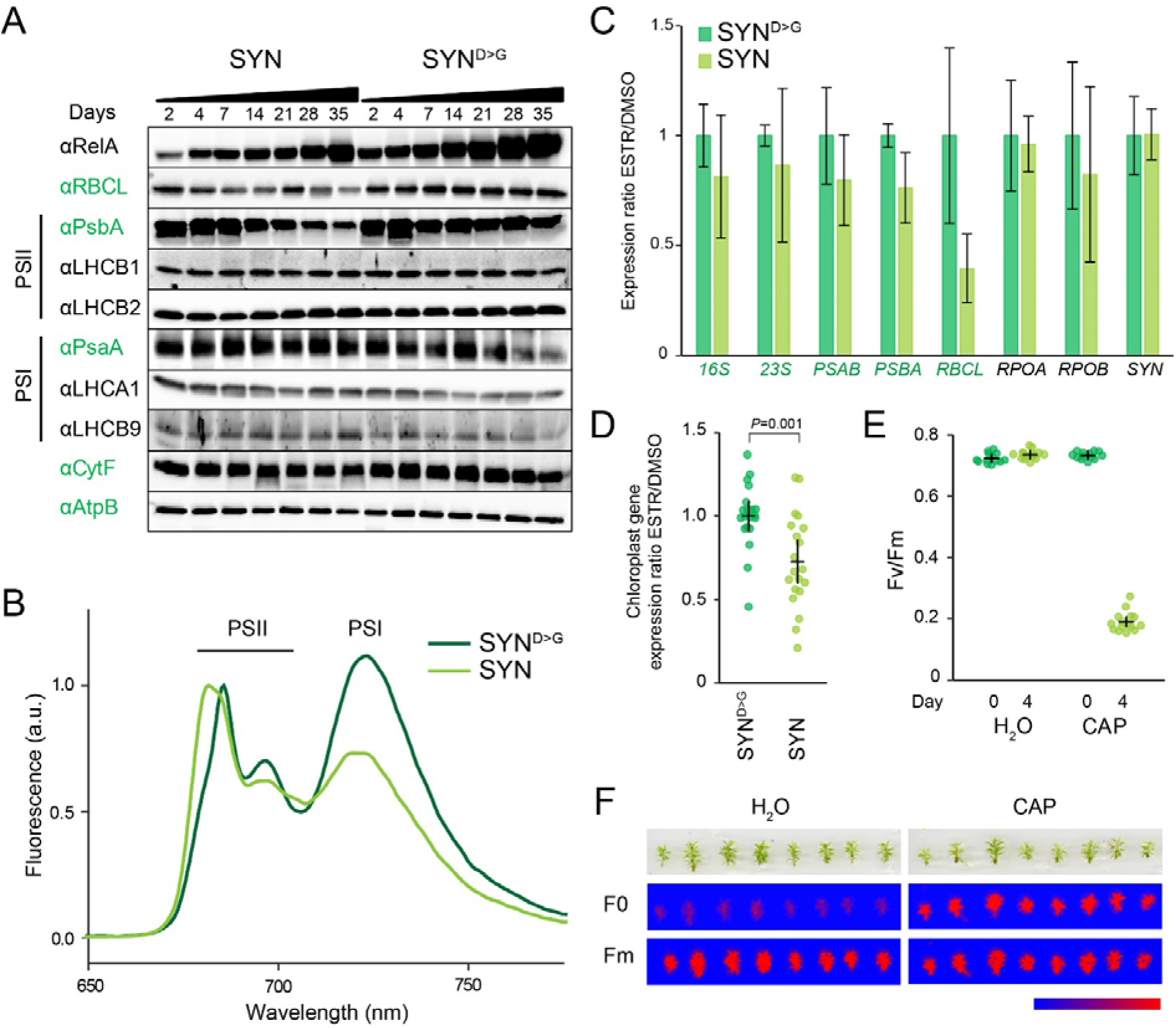
ppGpp accumulation alters chloroplast gene expression and reduces LHCII connectivity to PSII core. (A) Immunoblots on equal quantities of total protein from SYN and SYN^D>G^ estradiol induced gametophores using antibodies against signature chloroplast and nuclear encoded photosynthetic proteins. Chloroplast-encoded proteins are indicated by green text. (B) Low temperature (77K) fluorescence emission spectra of SYN and SYN^D>G^ gametophores. Spectra are normalized to maximum PSII emission. (C) RT-qPCR for selected chloroplast-encoded transcripts from SYN and SYN^D>G^ gametophores after 35 DOI (n= 4 biological replicates). (D) Comparison of the chloroplast gene expression ratio ESTR/DMSO in SYN and SYN^D>G^ for the chloroplast genes in (B), *P* calculated by two-way Student t-test. The effects of the chloroplast translation inhibitor chloramphenicol (CAP) on (E) the maximal efficiency of PSII (Fv/Fm) four days after treatment with CAP and (F) basal (F0) and maximal (Fm) chlorophyll fluorescence (false color scale, 20-1000 intensity units). Error bars indicate 95% CI.

The increase in the ratio of PSII light harvesting proteins (LHCII) to PSII RC (Fig 2A, Fig. S5) observed in induced SYN gametophores may contribute to the increased chlorophyll fluorescence (F0) by reducing the efficiency of energy transfer from LHCII into photochemistry. To test this hypothesis, we measured steady state chlorophyll fluorescence at 77K in SYN and SYN^D>G^ gametophores (Fig. 2B). In SYN^D>G^ the expected PSI fluorescence peak at 720 nm and PSII peaks at 685 nm and 695 nm were observed. In the SYN line the first PSII peak showed a striking blueshift to 681 nm and an increase in intensity with respect to PSI. The chlorophylls in isolated LHCII trimers fluoresce at 680 nm, while in intact PSII complexes the fluorescence shifts to 685 nm characteristic of the coupling between LHCII and the low energy chlorophylls of the PSII RC (Lamb *et al*., 2018). These results indicate that a substantial proportion of the excess LHCII trimers in induced SYN gametophores are energetically uncoupled from the PSII RC, and explains the increase in F0 and decrease in Fv/Fm in induced SYN gametophores.

In *A. thaliana*, ppGpp affects chloroplast gene expression by inhibiting the transcription of chloroplast genes, and in particular by inhibiting the accumulation of chloroplast ribosomal RNA (Yamburenko *et al*., 2015; Sugliani *et al*., 2016). We therefore analysed the effects of ppGpp accumulation on chloroplast gene expression by RT-qPCR (Fig. 2C). SYN induction was associated with a small decrease in the abundance of the transcripts for all the chloroplast genes tested, with the most substantial decrease observed for *RBCL*. While the differences were not significant for individual genes, there was overall a clear and significant decrease in the average expression ratio of chloroplast genes in SYN following induction (Fig. 2D).

We next asked whether the inhibition of chloroplast gene expression alone was sufficient to explain the changes in photosynthesis induced by ppGpp accumulation. We applied chloramphenicol, an antibiotic that inhibits plastid gene expression, to wild-type gametophores and monitored its effect on photosynthetic capacity (Fig. 2E, F). After four days we observed a large rise in F0, and a corresponding drop in Fv/Fm similar to that which occurs following SYN induction (Fig. 1E). This indicates that inhibition of plastid gene expression by ppGpp is sufficient to explain the changes we observe in phostosystem organisation and activity.

### ppGpp accumulation causes major changes in chloroplast structure

In *A. thaliana*, ppGpp accumulation causes a reduction in chloroplast volume that is partially compensated for by an increase in chloroplast number (Sugliani *et al*., 2016). We therefore visualized chloroplasts in the phyllids of SYN and SYN^D>G^ lines by microscopy. Induced SYN chloroplasts appeared slightly smaller than chloroplasts in SYN^D>G^ (Fig. 3A and Fig. S6) or in the non-induced control, but there was no obvious difference in chloroplast number. However, to our surprise the chloroplasts in the induced SYN line contained dense round structures (8.56 ± 0.40 per chloroplast, n= 75 chloroplasts; diameter 2.09 μm ± 0.06, n= 68) that were absent from the controls (Fig. 3A, B). The dense round structures were present in the chloroplasts of all the SYN lines (Fig. S6), and started to form three to four days after induction, becoming progressively larger and more well defined with time (Fig. S7).

**Figure 3.**
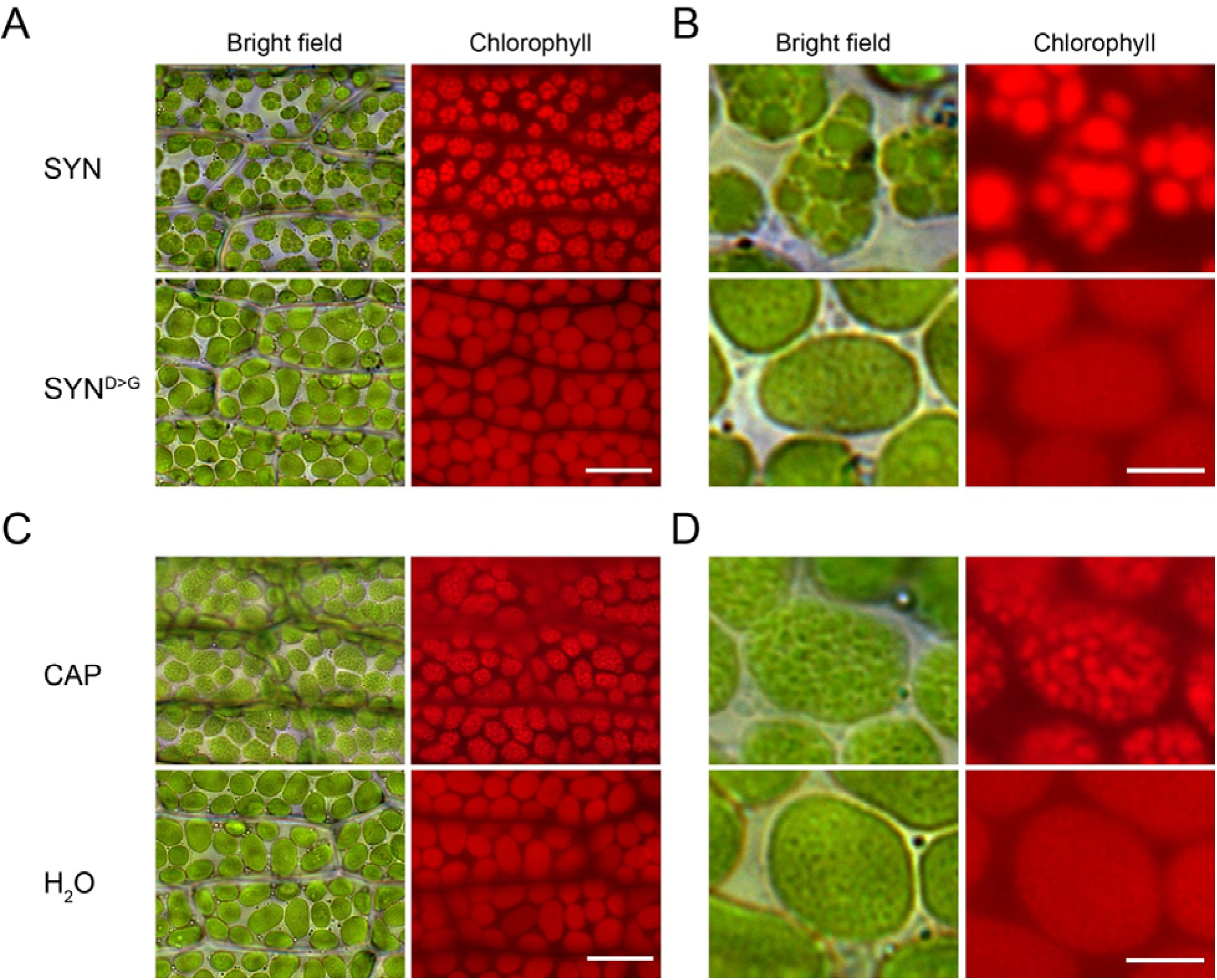
ppGpp accumulation causes a major reorganization of chloroplast structure. (A) Bright field and fluorescence microscopy images of phyllids from SYN1 and SYN^D>G^ gametophores at 35 DOI (scale, 20 μm). (B) Close-up images of single chloroplasts in A (scale, 5 μM). (C) Images of phyllids from wild-type gametophos7res 4 days after treatment with chloramphenicol (CAP) (scale, 20 μm). (D) Close-up images of single chloroplasts in C (scale, 5 μM).

We also found that the inhibition of chloroplast gene expression with chloramphenicol was sufficient to cause the appearance of similar dense round structures (Fig. 3C, D), as we also observed for chlorophyll fluorescence (Fig. 2E, F). Together, these results indicate that ppGpp accumulation drives a major structural change in *P. patens* chloroplasts that is associated with alterations in photosynthetic capacity and which is likely to be the result of the inhibition of chloroplast gene expression.

### ppGpp accumulation causes the formation of super grana

We used transmission electron microscopy to determine the nature of the structures in the chloroplasts of induced SYN and SYN^D>G^ gametophores (Fig. 4A). SYN^D>G^ chloroplasts contained clear stromal thylakoid membranes with well-defined grana. In contrast SYN chloroplasts contained only two to three very large grana structures (super grana) along the full length of the chloroplast.

**Figure 4.**
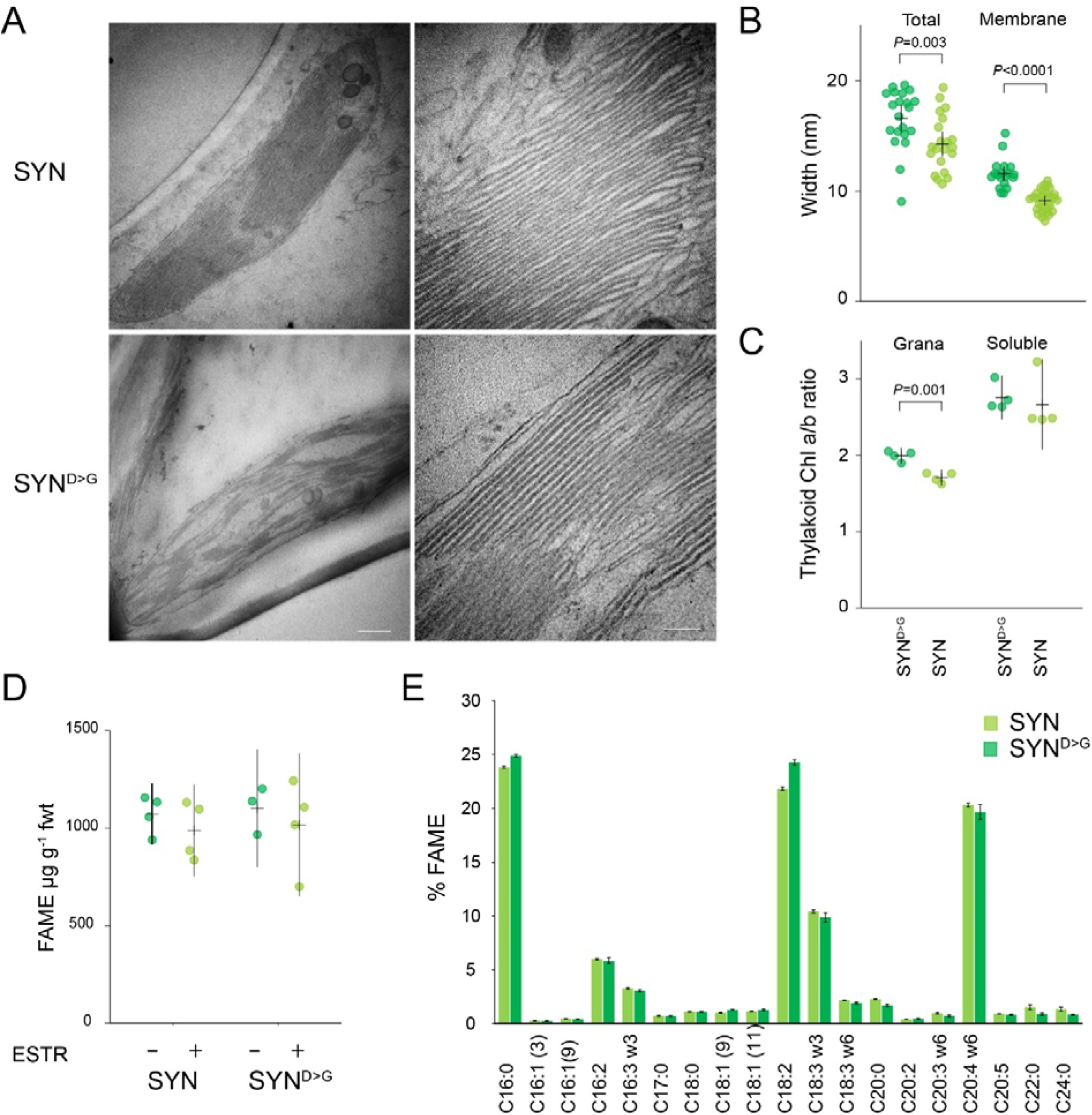
ppGpp accumulation causes the formation of super grana. (A) Electron microscopy images of SYN and SYN^D>G^ chloroplasts from gametophores after 35 DOI (scale 500 nm left panels, 100 nm right panels). (B) Average stack and membrane width in SYN and SYN^D>G^ grana (n=20 membranes, two-way Student t-test). (C) Chlorophyll a/b ratios for grana and soluble thylakoid fractions (n=3 biological repeats, two-way Student t-test). Fatty acid methyl ester (FAME) content (D) and composition (E) in SYN and SYN^D>G^ gametophores after 35 DOI (n=4 biological repeats). Measurments from ESTR induced gametophores are shown in (E). Error bars indicate 95% CI.

The super grana traversed the complete width of the chloroplast, and were separated by disorganised stromal lamellae. The thylakoid membranes in the SYN super grana were stacked more tightly than in SYN^D>G^, with reduced membrane and total repeat width (Fig. 4B). The super grana also showed a strikingly low chlorophyll a/b ratio compared to regular grana (Fig. 4C). LHCII trimers have a low chlorophyll a/b ratio of around 1.4 while the PSII RC and PSI have higher chlorophyll a/b ratios (Pinnola *et al*., 2015). These data indicate that SYN chloroplasts not only have more grana volume than in the control line, but that the grana membranes themselves are also more highly enriched in LHCII trimers.

The super grana in induced SYN gametophores represent a major reorganisation of the chloroplast membrane system. We therefore suspected that the super grana could be the result of membrane lipid synthesis or remodeling. However, we found no significant difference in fatty acid content (Fig. 4D) and only minor changes in fatty acid composition (Fig. 4E) indicating that de novo lipid biosynthesis is unlikely to play a major role in super grana formation. Furthermore, analysis of the composition of polar lipid classes also revealed no major changes in the galactolipids monogalactosyldiacylglycerol (MGDG) and digalactosyldiacylglycerol (DGDG) that are the main components of chloroplast membranes (Fig. S8).

## Discussion

In this study we examined the role of ppGpp in the moss *P. patens*. We created an inducible expression system to trigger the accumulation of ppGpp (SYN), with controls that allowed us to separate the specific effects of ppGpp from other effects of the inducible expression system (SYN^D>G^) (Fig.1, Fig. S1). Using these lines we were able to explore the likely role of ppGpp in *P. patens*, and to make comparisons with the role of ppGpp signalling in other photosynthetic organisms.

The data we present indicate that the effect of ppGpp on photosynthetic function and in particular on PSII is highly conserved across plants and algae. We found that ppGpp accumulation in gametophores caused a major decrease in photosynthetic capacity as shown by a reduction in multiple photosynthetic parameters and the levels of several chloroplast encoded proteins including Rubisco (Fig. 1–2). Similar, although less pronounced, photosynthetic phenotypes were also reported in *A. thaliana* plants that overaccumulate ppGpp (Maekawa, M. *et al*., 2015; Sugliani *et al*., 2016) as well as more recently in the diatom *P. tricornutum* (Avilan *et al*., 2020). The conservation of core ppGpp responsive elements is in line with the early origins of the RSH enzyme family for ppGpp homeostasis in the photosynthetic eukaryotes (Atkinson *et al*., 2011; Ito *et al*., 2017; Avilan *et al*., 2019).

Our results in *P. patens* support the hypothesis that ppGpp acts principally via the inhibition of chloroplast gene expression in plants. In *A. thaliana* ppGpp inhibits chloroplast gene expression via the inhibition of chloroplast transcription, although the molecular mechanism is not yet known (Sugliani *et al*., 2016; Field, 2018). Here we found that ppGpp accumulation caused a small but significant decrease in the transcript levels of chloroplast encoded genes, consistent with a reduced rate of transcription (Fig. 2). The accumulation of a number of chloroplast encoded proteins, in particular PsbA and RBCL, was also inhibited by ppGpp accumulation (Fig. 2A). However, we note that some chloroplast-encoded proteins such as PsaA appeared to increase in abundance, suggesting that other layers of regulation are also be at play. Nevertheless, in support of a general effect of ppGpp on chloroplast gene expression we also show that treatment with the chloroplast translation inhibitor chloramphenicol is sufficient to cause a similar photosynthetic phenotype to ppGpp accumulation (Fig. 2).

Certain aspects of the response to ppGpp accumulation in *P. patens* were different to those observed in other organisms. In *A. thaliana* plants ppGpp accumulation causes visible yellowing with a clear reduction in chlorophyll levels (Maekawa, M. *et al*., 2015; Sugliani *et al*., 2016). However, here in *P. patens* we observed no detectable effect on gametophore chlorophyll in response to ppGpp accumulation under standard growth conditions (Fig. 1). While we observed a relatively small fold increase in ppGpp levels in response to SYN induction in *P. patens* we also observed much stronger decreases in Fv/Fm and the chlorophyll a/b ratio than previously observed in *A. thaliana* (Sugliani *et al*., 2016), and we also observed the major reorganisation of chloroplast structure (Fig. 3–4). Therefore, we consider it likely that the different response of *P. patens* may reflect differences in the underlying modes of action for ppGpp rather than in the quantity of ppGpp. As such our findings may be evidence of the specialisation of ppGpp signalling in different branches of the photosynthetic eukaryotes.

We observed the formation of striking super grana structures in *P. patens* chloroplasts in response to ppGpp accumulation (Fig. 3–4). This was surprising because super grana were not previously observed in *A. thaliana* plants that overaccumulate ppGpp (Maekawa, M. *et al*., 2015; Sugliani *et al*., 2016). Super grana occur naturally in shade-adapted plants like *Alocasia macrorrhiza* (Heitz, 1936; Anderson, 1999), and can also be induced in some plants when exposed to low light levels such as in apple (Skene, 1974). *A. thaliana* does not accumulate super grana under low light. However, growth of *A. thaliana* on the chloroplast translation inhibitor lincomycin provokes the accumulation of super grana in leaves (Belgio *et al*., 2015). Interestingly, the lincomycin-grown plants have an Fv/Fm of around 0.2, much lower than the Fv/Fm observed in response to ppGpp over accumulation in *A. thaliana* and more similar to that observed in *P. patens* (Fig. 1). Interestingly, our data indicate that the reduced Fv/Fm and the accumulation of super grana are likely not due to changes in lipid composition (Fig. 4E, S8), but rather modifications in protein composition. The reduction in Fv/Fm in *P. patens* overaccumulating ppGpp and in lincomycin-grown *A. thaliana* is likely to be due to a substantial energetic uncoupling of PSII antenna from PSII reaction centres (Fig. 2B) (Belgio *et al*., 2015). One way that this can occur is by an increase in the proportion of PSII LHCII trimers to PSII reaction centres. Increased LHCII trimer abundance is also known to promote grana stacking (Pribil *et al*., 2014; Albanese *et al*., 2020). However, rather counter intuitively, overaccumulation of ppGpp resulted in an apparently higher LHCII trimer to PSII RC ratio in *A. thaliana* (Sugliani *et al*., 2016) than we oberve here in *P. patens* (Fig. 2A). An explication for this discrepancy is that photosystem composition in *P. patens* is different to that of *A. thaliana* (Alboresi *et al*., 2008), leading to uncertainty about the quantity and function of each component. For example, *P. patens* possesses many more LHCII isoforms, including the algal-like LHCB9 isoform that is absent from *A. thaliana*. Indeed, we noticed that LHCB9 accumulated in response to ppGpp (Fig. 2 and Fig S5). The increase in the abundance of this isoform, and potentially other isoforms for which we do not have specific antibodies could be responsible for promoting grana stacking and/or causing a larger change in the LHCII trimer / PSII RC ratio than we could observe by immunoblotting (Fig. 2A). This latter possibility is supported by the pronounced blue-shift in the low temperature chlorophyll fluorescence profile for SYN, indicating energetic uncoupling of LHCII trimers from PSII RC (Fig. 2B), as well as the low chlorophyll a/b ratio in gametophores and isolated grana (Fig. 1H, 4C).

Despite having a dramatic impact on thylakoid organisation and photosynthetic capacity, ppGpp accumulation did not alter gametophore growth (Fig. 1). Our results contrast with the situation in *A. thaliana* plants where overexpression of RSH3 or an equivalent SYN protein increased ppGpp levels, reduced photosynthetic capacity and caused a strong reduction in growth (Sugliani *et al*., 2016). Similarly, the induction of ppGpp synthesis by SYN in the diatom *P. tricornutum* reduced photosynthetic capacity while strongly repressing proliferation (Avilan *et al*., 2020). These observations strongly suggest that ppGpp has different impacts on growth in divergent photosynthetic eukaryotes. More generally, these results may reflect different responses to a reduction in photosynthetic capacity. Analysis of gas-exchange data shows that bryophytes obtain lower photosynthetic benefits on a per leaf mass area or N content basis than angiosperms (Gago *et al*., 2019). Gago et al. (2019) propose that this difference is related to the limitation of CO2 diffusion by the bryophyte cell wall, differential nutrient investment strategies, and a fundamentally different canopy structure to angiosperms. While it is not clear which of these factors is important, it is likely that they contribute to the low sensitivity of *P. patens* growth to the severe drop in photosynthetic rate and disruption of photosystem organisation caused by ppGpp accumulation. Comparison of the impact of these changes on stress tolerance in both species would need more investigation (Maekawa, M. *et al*., 2015; Honoki *et al*., 2018; Gago *et al*., 2019). It would also be interesting to investigate gametophore growth in *P. patens* mutants affected in photosystem composition to understand whether our observation is specific to ppGpp accumulation or reflects a more general ability of mosses to grow despite defects in photosynthesis (Pinnola *et al*., 2015; Pinnola *et al*., 2018; Peng *et al*., 2019; Storti *et al*., 2020). Analysis in other model bryophytes, such as the liverwort *Marchantia polymorpha*, would also help to determine whether this characteristic can be generalized to all non-vascular plants (Cesarino *et al*., 2020).

Altogether, our data indicate that a relatively modest increase in ppGpp can inhibit chloroplast gene expression in *P. patens* and lead to a major downregulation of photosynthetic activity and radical reorganisation of the chloroplast membrane system without a major impact on growth. Our data point to the existence of both highly conserved targets of ppGpp signalling as well as more specialised targets in different organisms. This may be related to the different relationship between photosynthetic capacity and growth observed between bryophytes and vascular plants. Further research is now required to understand the physiological roles of ppGpp during growth and development across land plants, and also the significance and mechanism of super grana formation.

## Acknowledgements

This work was supported by a CEA IRTELIS PhD fellowship for S.H., an APEX grant from the Provence-Alpes Cote d’Azur Region, and Agence Nationale de la Recherche funding (ANR-15-CE05-0021-03, SignauxBioNRJ). Nucleotide measurements were performed on the IJPB Plant Observatory technological platform which is supported by Saclay Plant Sciences-SPS (ANR-17-EUR-0007), microscopy experiments were performed on the PiCSL-FBI core facilty member of the France-BioImaging national research infrastructure (ANR-10-INBS-04), and lipidomics experiments were performed on the Heliobiotec platform (CEA Cadarache). We thank Julia Bartoli and Emmanuelle Bouveret for kindly providing 13C-ppGpp, Marie-Helene Montané for setting up moss culture methods, and Aurelie Crepin and Stefano Caffarri for assistance with grana isolation and low temperature chlorophyll fluorescence measurements.

## Contributions

B.M. and B.F. conceived the experiments. S.C. performed the nucleotide quantification; B.L. and Y.L-B. performed and analysed the lipidomics experiments; S.H and A.A performed the electron microscopy experiments; S.H., S.E. and J.V. performed the remaining experiments. S.H., S.E., B.M. and B.F contributed to the interpretation of the results. S.H., B.M. and B.F. wrote the manuscript. All authors provided critical feedback and helped shape the research, analysis and manuscript.

## Materials and Methods

### Plant growth

*P. patens* (Gransden strain) was grown under a 16h light/8h dark photoperiod at 23°C with 50 μmol m^−2^ s^−1^ PAR fluorescent lighting. For gametophore generation, fresh protonema cultivated on BCDAT agar media (Roberts *et al*., 2011) with cellophane disks was collected and dispersed in water with a T10 Basic ULTRA-TURRAX (IKA, Belgium) mixer, 3 ml of the homogenate were inoculated onto a sterile 7.4 mm diameter peat pellet (Jiffy products) and cultured for 25 days unless otherwise stated. To induce the expression of SYN, 25 day old peat pellet-moss cultures containing mainly young gametophores (Fig. S2) were treated with 5μM of estradiol (from stock dissolved in DMSO) or DMSO control two times a week for 35 days unless stated otherwise. For chloramphenicol tratments, 25 days old *P. patens* wild type cultures were sprayed once with 5 mM chloramphenicol (dissolved in water) or water control and then cultured under standard growth conditions.

### Cloning

A fragment corresponding to amino acids 1 to 386 of RelA was fused by PCR to a genomic sequence coding for the 80-amino acid *P. patens* Rubisco small subunit (Pp3c13_15980V3.1) target peptide. The fused PCR product (SYN) was then introduced into the entry vector pDONR207 (Life Technologies) by Gateway BP cloning. The entry clone was confirmed by sequencing. In parallel an inactive version of RelA with the mutation D275G was used in the same fashion to make SYN^D275G^.SYN and SYN^D275G^ were then recombined by LR Gateway recombination into the estradiol inducible expression vector pGGW6 (kindly provided by M. Hasebe) (Kubo *et al*., 2013).

### *P. patens* transformation

Polyethylene glycol–mediated transformation of the wild-type strain of *P. patens* with linearized plasmid was performed as described previously (Schaefer & Zrÿd, 1997) with small modifications (Menand *et al*., 2007). Selection of stable transformants was performed on BCDAT medium containing 25 μg/ml hygromycin. Southern blot analysis was performed using DIG labelled probes according to the manufacturer’s instructions (Roche). Three SYN and three SYN^D>G^ lines were shown to have one insertion in the correct PIG locus and therefore were selected for the study (Fig. S1). All biochemistry experiments were done with SYN1 and SYN^D>G^2 lines.

### Chlorophyll fluorescence measurements

Gametophores were aligned in a petri dish and adapted to the dark for 20 min and then imaged in a Fluorcam FC 800-O imaging fluorometer (Photon System Instruments). F0 and Fm were imaged using the following settings (Shutter=3, Sensitivity=75, Super=50), with F0 acquired over a 4 s period. PSII maximum quantum yield was calculated as (Fm-F0)/Fm. For the relative electron transfer rate (ETR) plants were dark-adapted and exposed to 2 minutes periods of increasing actinic light intensity, with Fv’/Fm’ measurements at the end of each period. Relative ETR was calculated as Fv’/Fm’ X light intensity. Steady state 77K chlorophyll fluorescence measurements were obtained from gametophore powder suspended in 85% (w/v) glycerol, 10 mM HEPES, pH 7.5 as described previously (Galka *et al*., 2012).

### Immunoblotting

Proteins were extracted and equal quantities of total protein analyzed by immunoblotting as described previously (Sugliani *et al*., 2016). The following antibodies were used: AtpB (Agrisera; polyclonal, catalog numberAS08304), LHCA1 (Agrisera; polyclonal, catalog number AS01 005), LHCB1 (Agrisera; polyclonal, catalog number AS01 004), LHCB9 (Agrisera; polyclonal, catalog number AS15 3088), PetA (Agrisera; polyclonal, catalog number AS08 306), PsaA (Agrisera; polyclonal, catalog number AS04 042), PsbA (Agrisera; polyclonal, catalog number AS05 084, and RelA (1/2000 dilution, raised against *E. coli* RelA and kindly provided by M.Cashel).

### Gene expression analysis

RNA was extracted from frozen gametophore powder using TriReagent (Sigma-Aldrich), quality was verified by agarose gel electrophoresis, and genomic DNA was removed by treatment with DNase (Thermofisher). cDNA was synthesized from 500 ng of RNA using Primescript RT Reagent Kit (Takara Bio) with random hexamer primers. qRT-PCR was performed on 1 μL of 1 in 40 diluted cDNA in 15 μL reactions using SYBR Premix Ex-Taq II reagent (Takara Bio) in a Bio-Rad CFX96 real-time system (see Table S1 for primer pairs). Relative quantification of gene expression adjusted for efficiency was performed using PCR Miner (Zhao & Fernald, 2005). *E2, 60S* and *APRT* were used as reference transcripts for normalization (Le Bail *et al*., 2013).

### Chlorophyll quantification

Chlorophyll was extracted from gametophores with ice-cold 80% acetone saturated with sodium carbonate. The absorbance was measured between 350 and 750 nm in a Varian Cary 300 spectrophotometer (Agilent). Chlorophyll concentrations and chlorophyll a/b ratios were calculated using a fitting algorithm as described previously (Croce *et al*., 2002).

### Quantification of ppGpp

ppGpp was extracted from gametophores and quantified by LC-MS/MS using a 13C-labelled G4P internal standard as described previously (Bartoli *et al*., 2020).

### Light and fluorescence microscopy

Gametophores were harvested and fixed in a solution of 2% glutaraldehyde 0.1 M phosphate buffer and stored at 4 °C overnight. Chloroplast structure was visualized in dissected phyllids using an Axioimager M2 microscope and Axiocam HRc Camera (Carl Zeiss Microscopy, Marly le Roi, and France). Chlorophyll auto-fluorescence was visualized with HBO 100 mercury lamp with excitation at BP 560/55, and emission at BP 645/75. Images were captured with AxioVision Rel 4.8 software, and Zerene stacker software version 1.04 was used for focus stacking correction.

### Electron microscopy

Phyllids were cut from gametophores and fixed in 2.5 % glutaraldehyde. Phyllids were transferred into phosphate buffer pH 7.2 for 5 min, incubated 5 min in agarose at 37°C and left overnight at 4°C. Phyllids were then washed twice 15 min in phosphate buffer 7.2 pH, post-fixed in 1% Osmium phosphate buffer 7.2 pH, 30 min at room temperature, and washed twice in water. The phyllids were then taken through a series of dehydratation steps in acetone at 20, 35, 50, 62, 75, 85, & 95 % for 15 min each and 3 times 100% for 5 min. The phyllids were kept in acetone and taken through a series of infiltration with SPURR resin (Sigma) (5, 10, 20, 40, and 70 %) 30 min each without accelerator and then 100% with SPURR S2 for 2 hr, and changed to fresh resin and left overnight. The next day, the resin was changed and left for 24 hr. Next, samples were placed in a mould with fresh resin and incubated at 60°C for 48 hr. The blocks were cut into ultra-thin sections of 70 nm using a Cryo-ultramicrotome UCT (Leica). The sections were placed on an EM grid, stained with aqueous uranyl acetate and lead citrate and observed with a FEI Tecnai G2 electron microscope. ImageJ software version 1.45 was used to measure membrane thickness.

### Analysis of thylakoid composition

Thylakoid membrane preparation and solubilization was performed as described previously, with modifications (Berthold *et al*., 1981). Approximately 10 g of gametophore was blended in 50 ml of buffer B1 (0.4 M NaCl, 2 mM MgCl_2_, 20 mM tricine KOH pH 7.8, 0.2 mM benzamidine, 1 mM hexanoic acid) and passed through a 30 μM filter. The solution was centrifuged 15 min at 1400 g and 4°C. Pellets were re-suspended softly with a paint brush in buffer B2 (20 mM tricine KOH pH 7.8, 0.15 M NaCl, 5 mM MgCl_2_, 0.2 mM benzamidine, 1 mM hexanoic acid) and centrifuged for 10 min at 6000g and 4°C. The pellets representing total thylakoids were then re-suspended in buffer B3 (15 mM NaCl, 5 mM MgCl_2_, 20 mM Hepes KOH pH 7.5) at a chlorophyll concentration of 2.5 mg ml^−1^. For solubilisation 100 μL of total thylakoid was mixed with 18 μL of Triton buffer (15% (w/v) triton X-100, 15 mM NaCl, 5 mM MgCl2) and incubated 20 min in the dark on ice with gentle agitation. The mix was centrifuged 3 min at 1500 g and 4°C and then the supernatant was submitted to a final centrifugation for 30min at 29500 rpm at 4°C in an Optima LE-80k ultracentrifuge. The supernatant, enriched for soluble stromal membranes, were separated from the Triton resistant pellets enriched in grana and chlorophyll measured in each fraction.

### Lipid measurements

Lipids were extracted from gametophores after 35 DOI as described previously (Hara & Radin, 1978) with modifications.Two ml of pre-heated isopropanol (85°C) containing BHT 0.01% (w/v) was added to 200-400 mg frozen gametophore powder in a glass tube with Teflon lined cap. The mixture was vortexed to break the tissue and heated for 5-10 minutes at 85°C in a water bath. After cooling down, internal standards (1 μg PE17:0/17:0, 1 μg TAG17:0/17:0/17:0), and 6 ml of MTBE were added to the mixture and vortexed. To allow phase separation, 2 volumes of aqueous sodium chloride NaCl (0.9 w/v) were added and the solutions were vortexed and centrifuged at 3000 *g* for 2 min. The upper phase containing lipids was collected and transfered to a new glass tube. To maximize extraction, 1 ml of MTBE was added to the samples again and vigorously vortexed and centrifuged at 3000 g for 2 min. The new upper phase was transfered to the tube containing the first solvent extract. Finally, lipid extracts were stored at −20°C until analysis. Fatty acids were first converted to their methyl esters then analyzed by gas chromatography-mass spectrometry (GC-MS) and lipid molecular species were analyzed by ultra performance liquid chromatography - tandem mass spectrometer (UPLC-MS/MS) as previously described (Legeret *et al*., 2016)

## Supplementary information

**Figure S1.**
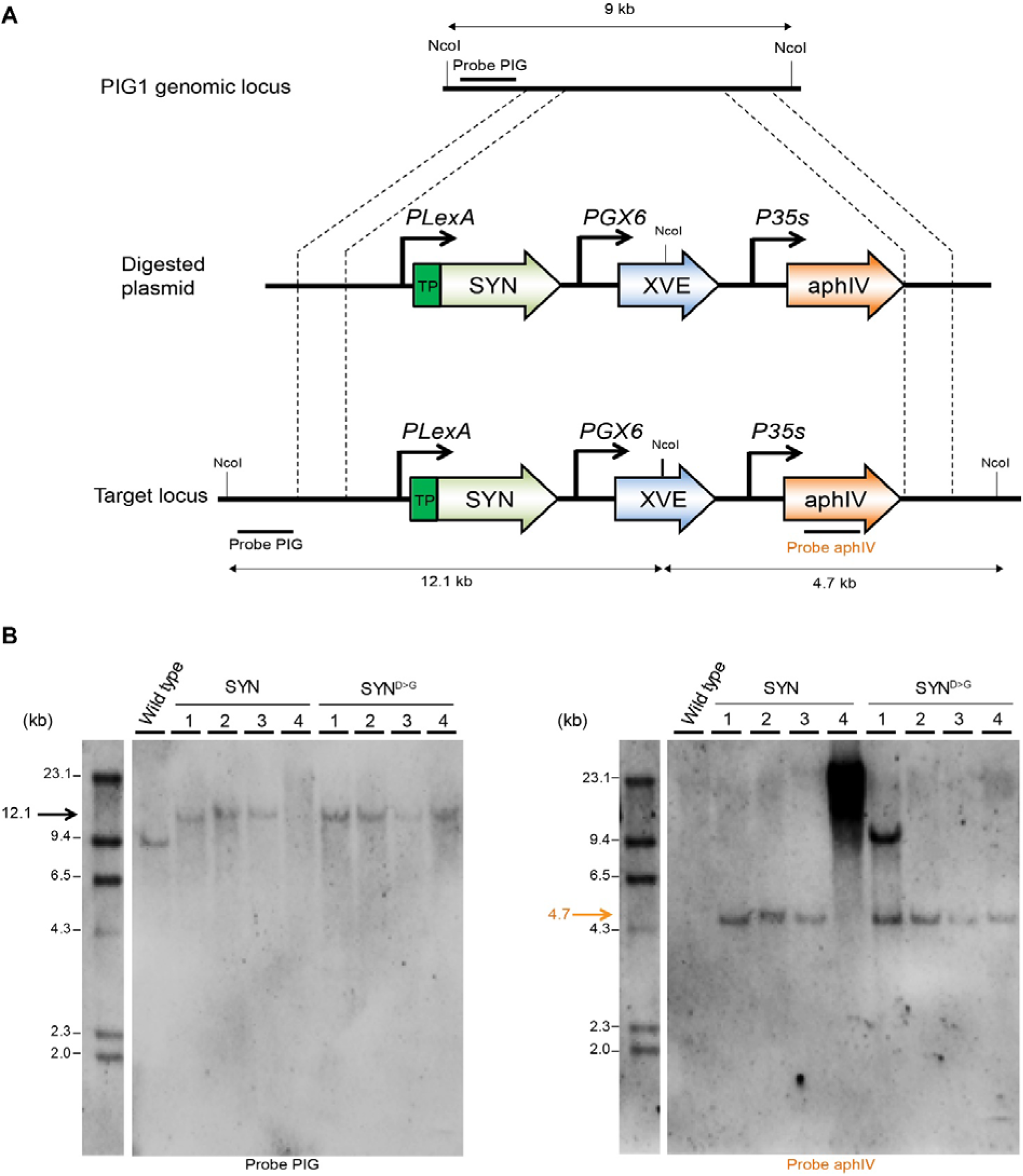
Southern blot analysis of *P. patens* transgenic SYN lines. (A) Schematic representation of the PIG locus and the gene targeting construct. Probes for Southern blotting and the expected size of the restriction fragments they should detect after successful gene targeting are indicated. TP: transfer peptide, SYN: ppGpp synthetase, XVE: chimeric transcription activator composed of: LexA (X), VP16 (V) and the human estrogen receptor (E), aphIV: hygromycin phosphotransferase resistant gene, PIG : *P. patens* Intergenic 1. (B) DNA extracted from different SYN lines was digested with *Nco*I and hybridized using a DIG labelled probe specific to the PIG locus (left), and then the same membrane was re-probed with a DIG-labelled probe specific to the antibiotic resistance cassette found in the gene targeting construct (right). The lines SYN 1, 2, 3 and SYN^D>G^ 2, 3, 4 were selected for further study.

**Figure S2.**
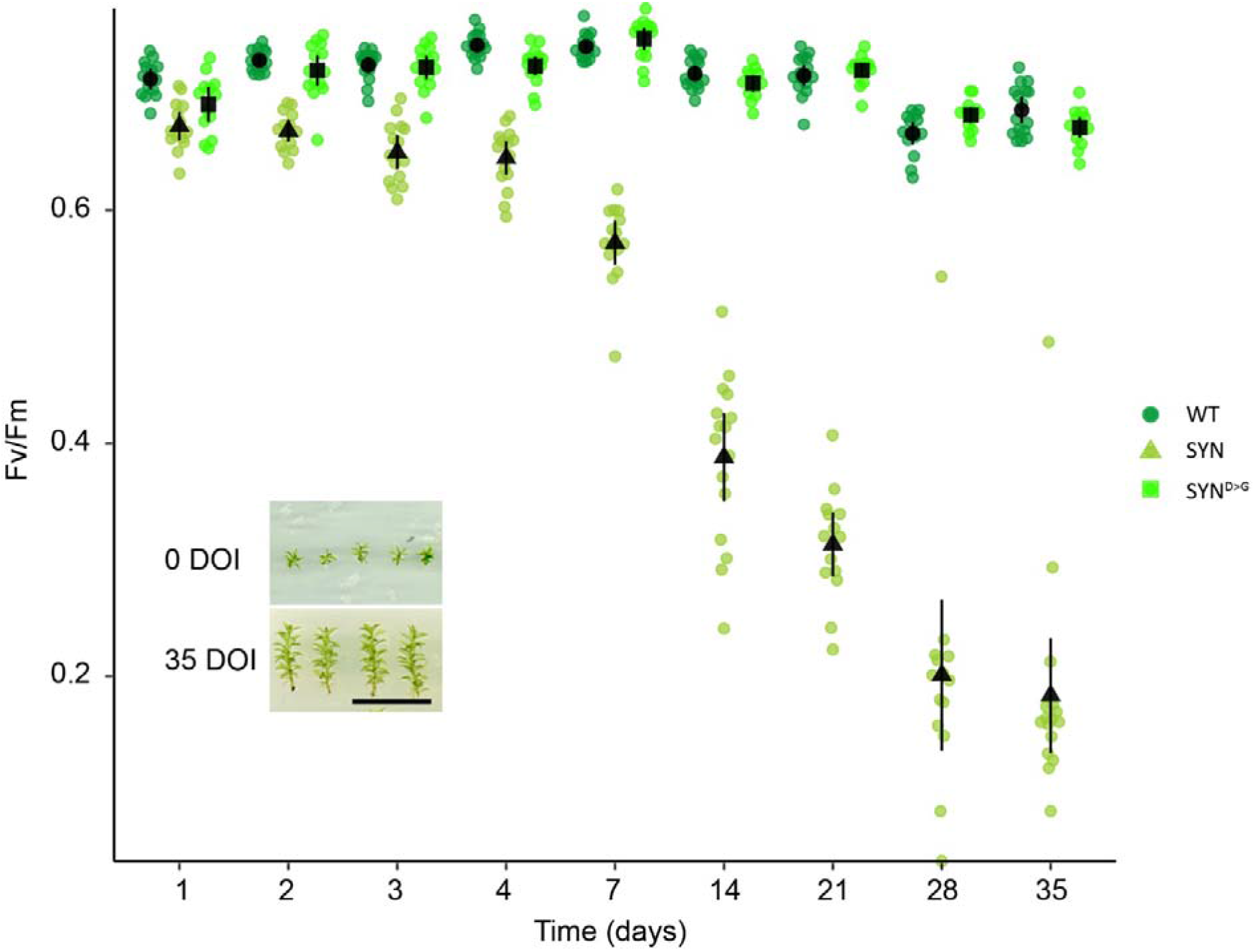
Evolution of Fv/Fm in different lines in response to estradiol treatment. Fv/Fm was measured regularly following the start of estradiol treatment in SYN, SYN^D>G^ and wild-type gametophores (n= 15 gametophores). Inset shows typical gametophore size at the start and end of the experiment (DOI, days of induction). Scale, 1cm. Error bars indicate 95% CI.

**Figure S3.**
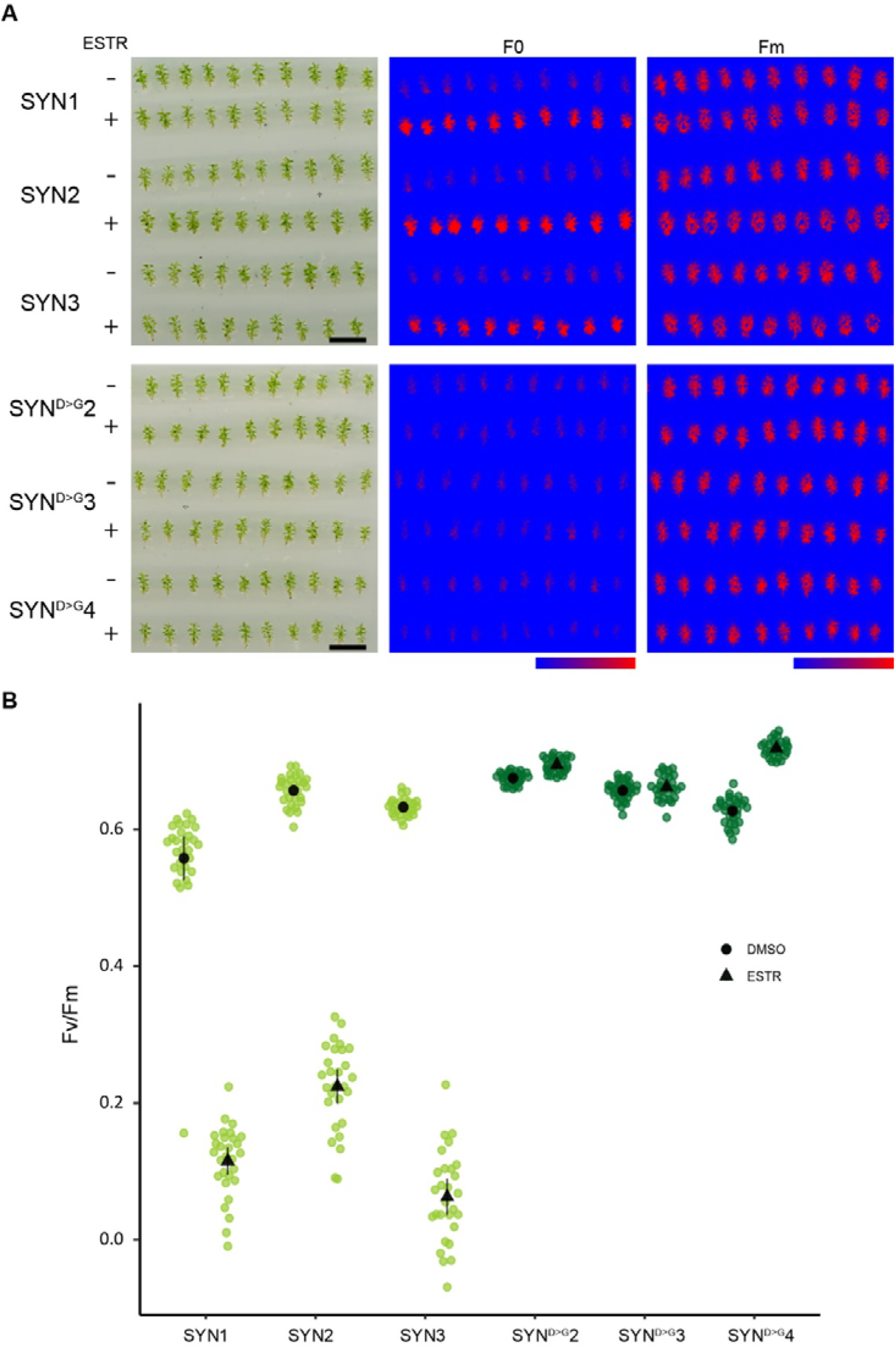
Photosynthetic parameters in different SYN lines. (A) Images of F0 and Fm in different lines (scale, 1 cm; false color scale, 50-2000 intensity units). (B) Fv/Fm in SYN and SYN^D>G^ lines after 35 DOI (n=30 gametophores). Error bars indicate 95% CI.

**Figure S4.**
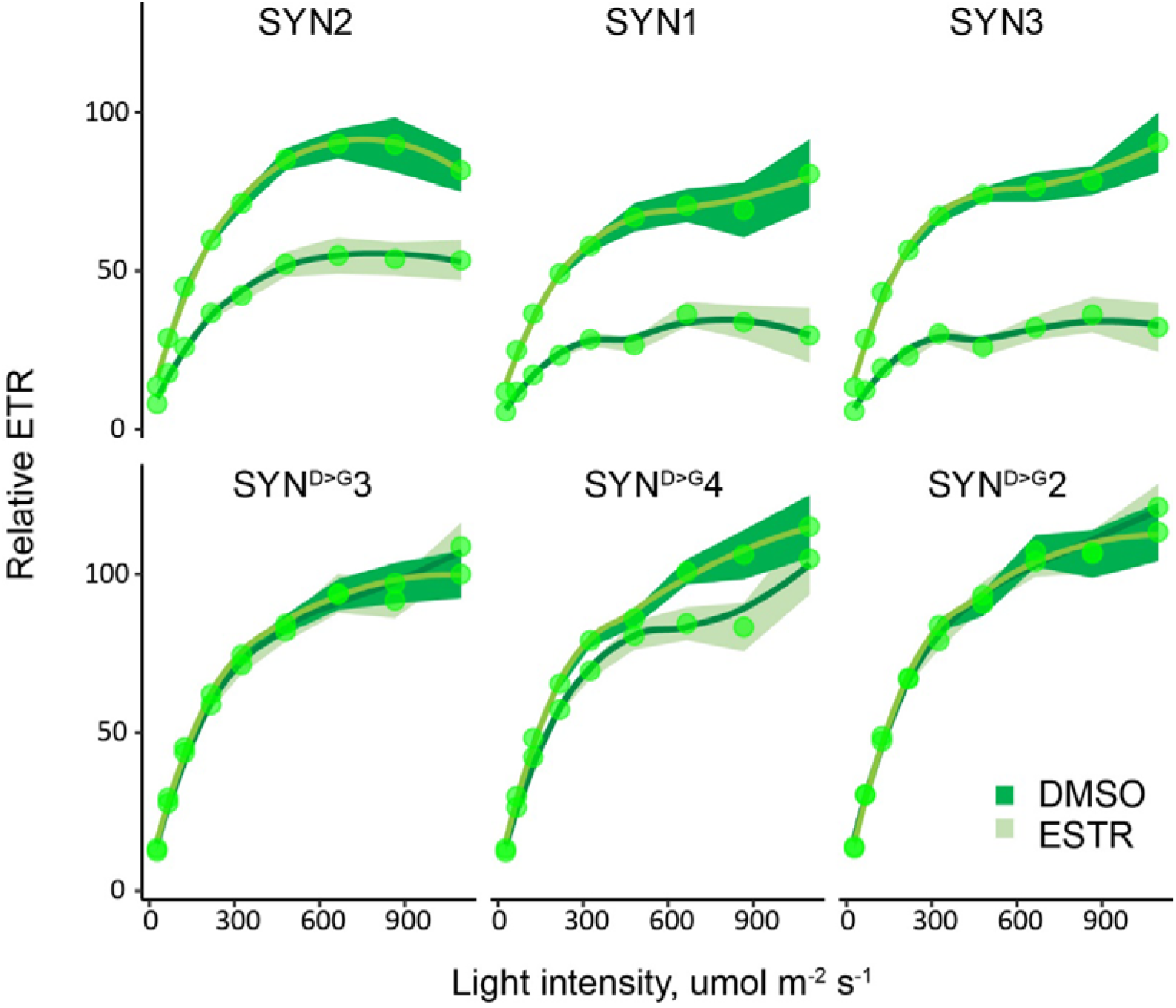
ETR in different SYN lines. ETR was determined at different light intensities in SYN and SYN^D>G^ gametophores after 35 DOI (n=30 gametophores). Error ribbon indicates 95% CI.

**Figure S5.**
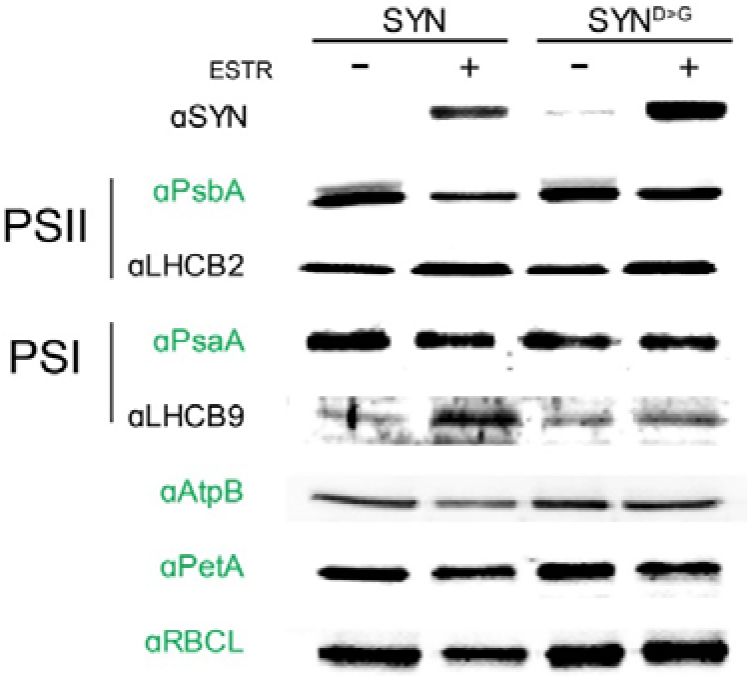
Protein accumulation after 35 DOI. Immunoblots on equal quantities of total protein from SYN and SYN^D>G^ after 35 DOI with 0 μM or 5μM estradiol using antibodies against signature chloroplast and nuclear encoded photosynthetic proteins. Chloroplast-encoded proteins are indicated by green text.

**Figure S6.**
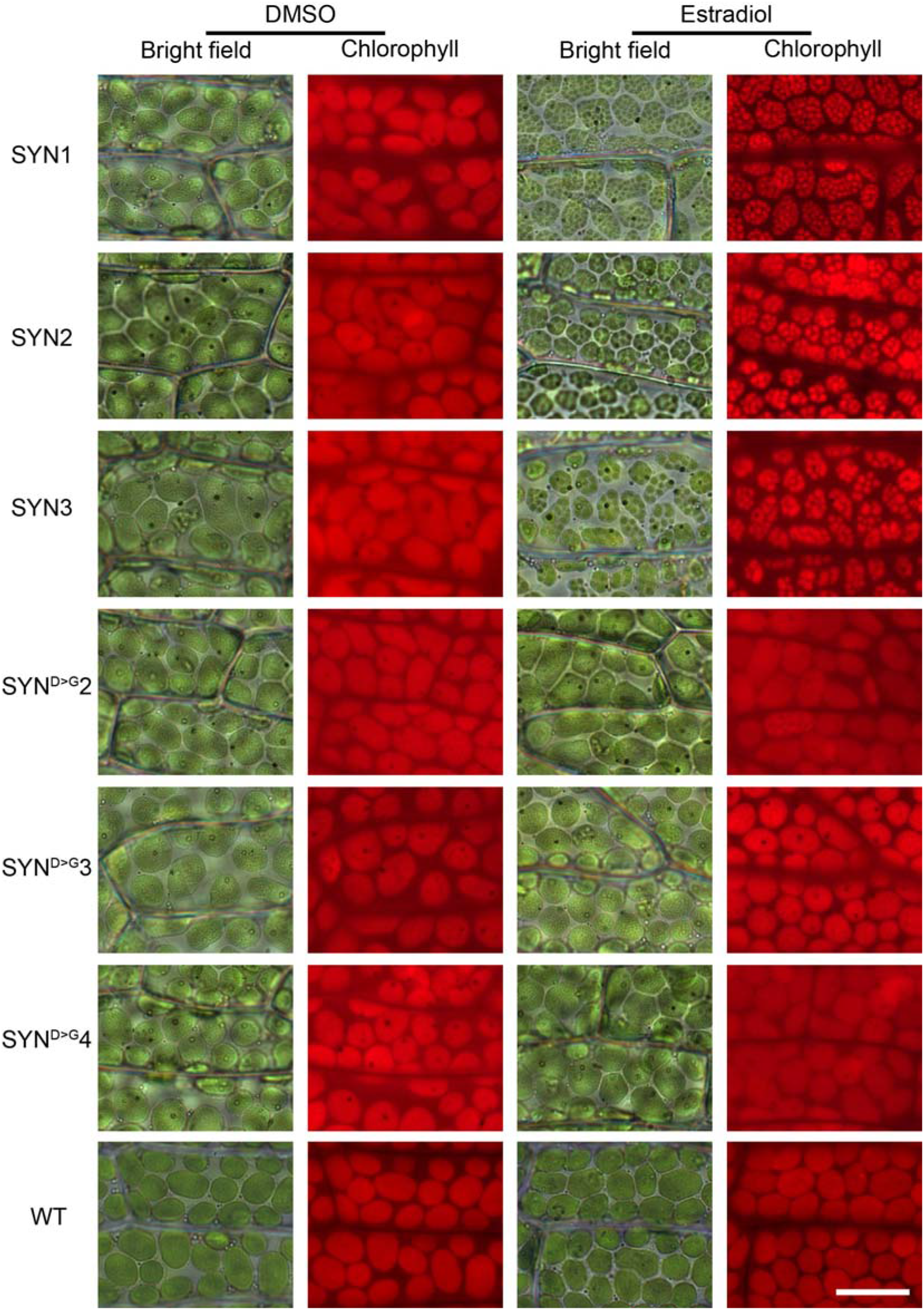
Chloroplast structure in independent SYN lines. Bright field and fluorescence microscopy images of phyllids from different SYN lines after 35 DOI (scale, 20 μM).

**Figure S7.**
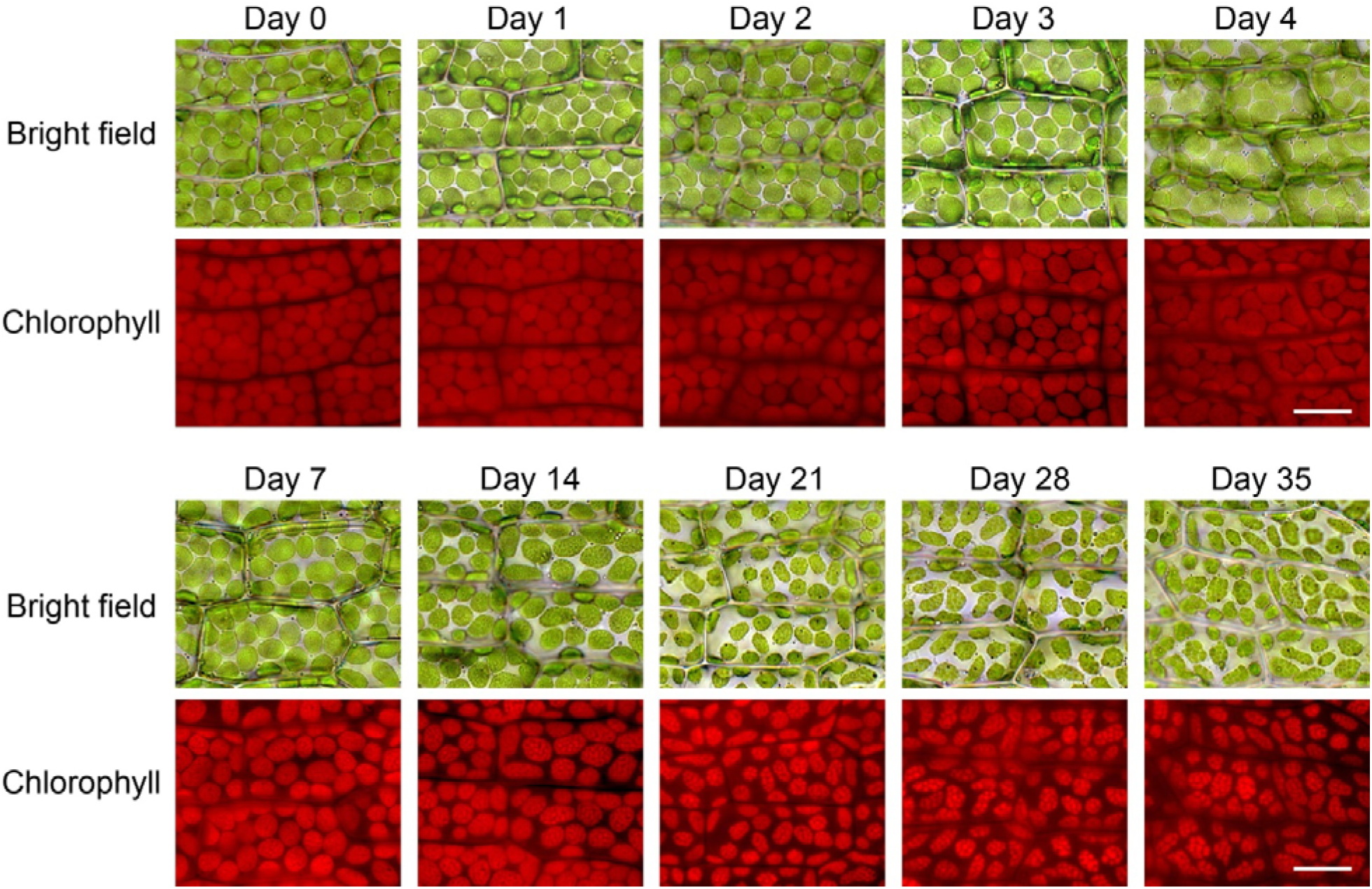
Appearance and evolution of super grana during SYN induction. Bright field and fluorescence microscopy images of phyllids at multiple timepoints during the induction of SYN (scale, 20 μM).

**Fig. S8.**
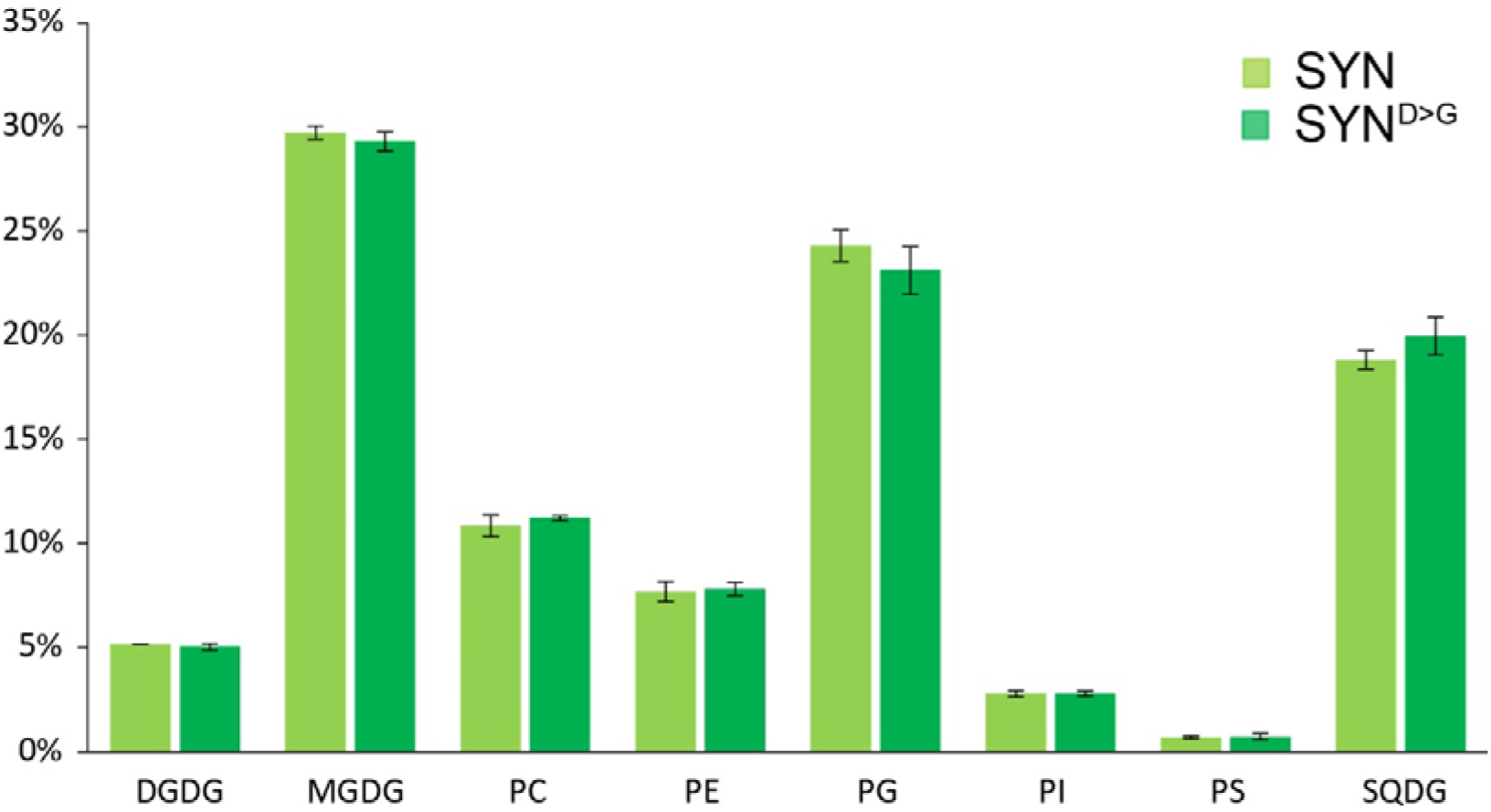
Polar lipid composition of induced SYN and SYN^D>G^ gametophores. Polar lipids were measured in gametophores 35 DOI (n=4 biological repeats). Error bars indicate 95% CI.

**Table S1.**
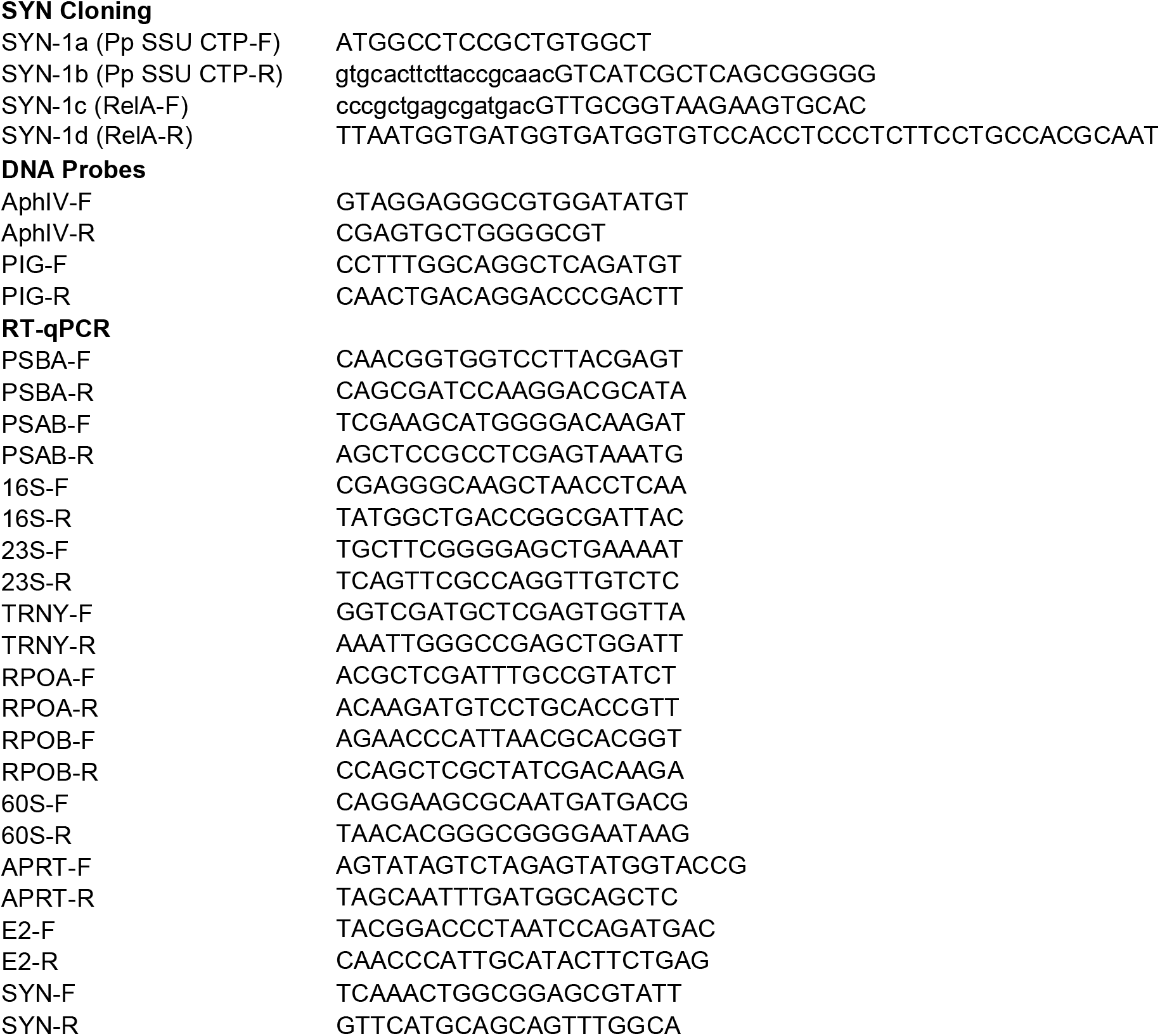
List of primers.

## References

Abdelkefi H, Sugliani M, Ke H, Harchouni S, Soubigou-Taconnat L, Citerne S, Mouille G, Fakhfakh H, Robaglia C, Field B. 2018. Guanosine tetraphosphate modulates salicylic acid signalling and the resistance of Arabidopsis thaliana to Turnip mosaic virus. Molecular Plant Pathology 19(3): 634–646.

Albanese P, Tamara S, Saracco G, Scheltema RA, Pagliano C. 2020. How paired PSII–LHCII supercomplexes mediate the stacking of plant thylakoid membranes unveiled by structural mass-spectrometry. Nature Communications 11(1): 1361.

Alboresi A, Caffarri S, Nogue F, Bassi R, Morosinotto T. 2008. In silico and biochemical analysis of Physcomitrella patens photosynthetic antenna: identification of subunits which evolved upon land adaptation. PLoS One 3(4): e2033.

Anderson JM. 1999. Insights into the consequences of grana stacking of thylakoid membranes in vascular plants: a personal perspective. Functional Plant Biology 26(7):625–639.

Atkinson GC, Tenson T, Hauryliuk V. 2011. The RelA/SpoT homolog (RSH) superfamily: distribution and functional evolution of ppGpp synthetases and hydrolases across the tree of life. PLoS One 6(8): e23479.

Avilan L, Lebrun R, Puppo C, Citerne S, Cuiné S, Li-Beisson Y, Menand B, Field B, Gontero B. 2020. ppGpp influences protein protection, growth and photosynthesis in Phaeodactylum tricornutum. bioRxiv: 2020.2003.2005.978130.

Avilan L, Puppo C, Villain A, Bouveret E, Menand B, Field B, Gontero B. 2019. RSH enzyme diversity for (p)ppGpp metabolism in Phaeodactylum tricornutum and other diatoms. Sci Rep 9(1): 17682.

Bartoli J, Citerne S, Mouille G, Bouveret E, Field B. 2020. Quantification of guanosine triphosphate and tetraphosphate in plants and algae using stable isotope-labelled internal standards. Talanta 219: 121261.

Belgio E, Ungerer P, Ruban AV. 2015. Light-harvesting superstructures of green plant chloroplasts lacking photosystems. Plant Cell Environ 38(10): 2035–2047.

Berthold DA, Babcock GT, Yocum CF. 1981. A Highly Resolved, Oxygen-Evolving Photosystem-Ii Preparation from Spinach Thylakoid Membranes - Electron-Paramagnetic-Res and Electron-Transport Properties. Febs Letters 134(2):231–234.

Cesarino I, Dello Ioio R, Kirschner GK, Ogden MS, Picard KL, Rast-Somssich MI, Somssich M. 2020. Plant science’s next top models. Annals of Botany 126(1): 1–23.

Croce R, Canino G, Ros F, Bassi R. 2002. Chromophore organization in the higher-plant photosystem II antenna protein CP26. Biochemistry 41(23): 7334–7343.

Field B. 2018. Green magic: regulation of the chloroplast stress response by (p)ppGpp in plants and algae. J Exp Bot 69(11):2797–2807.

Fujita T, Sakaguchi H, Hiwatashi Y, Wagstaff SJ, Ito M, Deguchi H, Sato T, Hasebe M. 2008. Convergent evolution of shoots in land plants: lack of auxin polar transport in moss shoots. Evolution & Development 10(2):176–186.

Gago J, Carriquí M, Nadal M, Clemente-Moreno MJ, Coopman RE, Fernie AR, Flexas J. 2019. Photosynthesis Optimized across Land Plant Phylogeny. Trends in Plant Science 24(10): 947–958.

Galka P, Santabarbara S, Khuong TT, Degand H, Morsomme P, Jennings RC, Boekema EJ, Caffarri S. 2012. Functional analyses of the plant photosystem I-light-harvesting complex II supercomplex reveal that light-harvesting complex II loosely bound to photosystem II is a very efficient antenna for photosystem I in state II. Plant Cell 24(7):2963–2978.

Glime JM 2020. Ecophysiology of Development: Gametophore Buds. In:Glime JM ed. Bryophyte Ecology..

Hara A, Radin NS. 1978. Lipid extraction of tissues with a low-toxicity solvent. Anal Biochem 90(1): 420–426.

Harrison CJ, Morris JL. 2018. The origin and early evolution of vascular plant shoots and leaves. Philosophical Transactions of the Royal Society B: Biological Sciences 373(1739): 20160496.

Hauryliuk V, Atkinson GC, Murakami KS, Tenson T, Gerdes K. 2015. Recent functional insights into the role of (p)ppGpp in bacterial physiology. Nat Rev Microbiol 13(5):298–309.

Heitz E. 1936. Untersuchungen Über den Bau der Plastiden. Planta 26(1):134–163.

Honoki R, Ono S, Oikawa A, Saito K, Masuda S. 2018. Significance of accumulation of the alarmone (p)ppGpp in chloroplasts for controlling photosynthesis and metabolite balance during nitrogen starvation in Arabidopsis. Photosynth Res 135(1-3): 299–308.

Ihara Y, Ohta H, Masuda S. 2015. A highly sensitive quantification method for the accumulation of alarmone ppGpp in Arabidopsis thaliana using UPLC-ESI-qMS/MS. J Plant Res 128(3):511–518.

Ito D, Ihara Y, Nishihara H, Masuda S. 2017. Phylogenetic analysis of proteins involved in the stringent response in plant cells. J Plant Res 130(4): 625–634.

Kubo M, Imai A, Nishiyama T, Ishikawa M, Sato Y, Kurata T, Hiwatashi Y, Reski R, Hasebe M. 2013. System for stable beta-estradiol-inducible gene expression in the moss Physcomitrella patens. PLoS One 8(9): e77356.

Lamb JJ, Røkke G, Hohmann-Marriott MF. 2018. Chlorophyll fluorescence emission spectroscopy of oxygenic organisms at 77 K. Photosynthetica 56(1):105–124.

Le Bail A, Scholz S, Kost B. 2013. Evaluation of reference genes for RT qPCR analyses of structure-specific and hormone regulated gene expression in Physcomitrella patens gametophytes. PLoS One 8(8): e70998.

Legeret B, Schulz-Raffelt M, Nguyen HM, Auroy P, Beisson F, Peltier G, Blanc G, Li-Beisson Y. 2016. Lipidomic and transcriptomic analyses of Chlamydomonas reinhardtii under heat stress unveil a direct route for the conversion of membrane lipids into storage lipids. Plant Cell Environ 39(4): 834–847.

Maekawa M, Honoki R, Ihara Y, Sato R, Oikawa A, Kanno Y, Ohta H, Seo M, Saito K, Masuda S. 2015. Impact of the plastidial stringent response in plant growth and stress responses. Nat Plants 1: 15167.

Maekawa M, Honoki R, Ihara Y, Sato R, Oikawa A, Kanno Y, Ohta H, Seo M, Saito K, Masuda S. 2015. Impact of the plastidial stringent response in plant growth and stress responses. Nature Plants 1: 15167.

Menand B, Yi K, Jouannic S, Hoffmann L, Ryan E, Linstead P, Schaefer DG, Dolan L. 2007. An ancient mechanism controls the development of cells with a rooting function in land plants. Science 316(5830): 1477–1480.

Okano Y, Aono N, Hiwatashi Y, Murata T, Nishiyama T, Ishikawa T, Kubo M, Hasebe M. 2009. A polycomb repressive complex 2 gene regulates apogamy and gives evolutionary insights into early land plant evolution. Proc Natl Acad Sci U S A 106(38): 16321–16326.

Peng X, Deng X, Tang X, Tan T, Zhang D, Liu B, Lin H. 2019. Involvement of Lhcb6 and Lhcb5 in Photosynthesis Regulation in Physcomitrella patens Response to Abiotic Stress. International journal of molecular sciences 20(15): 3665.

Pinnola A, Alboresi A, Nosek L, Semchonok D, Rameez A, Trotta A, Barozzi F, Kouřil R, Dall’Osto L, Aro EM, et al. 2018. A LHCB9-dependent photosystem I megacomplex induced under low light in Physcomitrella patens. Nat Plants 4(11): 910–919.

Pinnola A, Cazzaniga S, Alboresi A, Nevo R, Levin-Zaidman S, Reich Z, Bassi R. 2015. Light-Harvesting Complex Stress-Related Proteins Catalyze Excess Energy Dissipation in Both Photosystems of Physcomitrella patens. Plant Cell 27(11):3213–3227.

Pribil M, Labs M, Leister D. 2014. Structure and dynamics of thylakoids in land plants. Journal of Experimental Botany 65(8):1955–1972.

Rensing SA, Goffinet B, Meyberg R, Wu S-Z, Bezanilla M. 2020. The *Moss Physcomitrium* (*Physcomitrella*) *patens*: A Model Organism for Non-Seed Plants. The Plant Cell 32(5): 1361.

Roberts AW, Dimos CS, Budziszek MJ, Jr., Goss CA, Lai V. 2011. Knocking out the wall: protocols for gene targeting in Physcomitrella patens. Methods Mol Biol 715:273–290.

Ronneau S, Hallez R. 2019. Make and break the alarmone: regulation of (p)ppGpp synthetase/hydrolase enzymes in bacteria. FEMS Microbiol Rev 43(4): 389–400.

Sato M, Takahashi T, Ochi K, Matsuura H, Nabeta K, Takahashi K. 2015. Overexpression of RelA/SpoT homologs, PpRSH2a and PpRSH2b, induces the growth suppression of the moss Physcomitrella patens. Biosci Biotechnol Biochem 79(1):36–44.

Schaefer DG, Zrÿd J-P. 1997. Efficient gene targeting in the moss Physcomitrella patens. The Plant Journal 11(6):1195–1206.

Skene DS. 1974. Chloroplast structure in mature apple leaves grown under different levels of illumination and their response to changed illumination. Proceedings of the Royal Society of London. Series B. Biological Sciences 186(1082): 75–78.

Steinchen W, Bange G. 2016. The magic dance of the alarmones (p)ppGpp. Mol Microbiol 101(4):531–544.

Storti M, Puggioni MP, Segalla A, Morosinotto T, Alboresi A. 2020. The chloroplast NADH dehydrogenase-like complex influences the photosynthetic activity of the moss Physcomitrella patens. Journal of Experimental Botany 71(18): 5538–5548.

Sugliani M, Abdelkefi H, Ke H, Bouveret E, Robaglia C, Caffarri S, Field B. 2016. An Ancient Bacterial Signaling Pathway Regulates Chloroplast Function to Influence Growth and Development in Arabidopsis. Plant Cell 28(3):661–679.

Takahashi K, Kasai K, Ochi K. 2004. Identification of the bacterial alarmone guanosine 5’-diphosphate 3’-diphosphate (ppGpp) in plants. Proc. Natl. Acad. Sci. USA 101(12): 4320–4324.

Yamburenko MV, Zubo YO, Borner T. 2015. Abscisic acid affects transcription of chloroplast genes via protein phosphatase 2C-dependent activation of nuclear genes: repression by guanosine-3’-5’-bisdiphosphate and activation by sigma factor 5. Plant J 82(6):1030–1041.

Zhao S, Fernald RD. 2005. Comprehensive algorithm for quantitative real-time polymerase chain reaction. J Comput Biol 12(8):1047–1064.

